# Nlrc4 inflammasome is critical for host protection against flagellated *Salmonella*

**DOI:** 10.1101/2022.05.13.491918

**Authors:** Américo H. López-Yglesias, Chun-Chi Lu, Marvin A. Lai, Ellen K. Quarles, Xiaodan Zhao, Adeline M. Hajjar, Kelly D. Smith

**Affiliations:** Department of Laboratory Medicine and Pathology, University of Washington. Seattle, WA. USA. 98195; Department of Comparative Medicine, University of Washington. Seattle, WA. USA. 98195

## Abstract

*Salmonella enterica* serovar Typhimurium is a leading cause of gastroenteritis worldwide and a deadly pathogen in children, immunocompromised patients, and the elderly. *Salmonella* induces innate immune responses through the Nlrc4 inflammasome, which has been demonstrated to have distinct roles during systemic and mucosal detection of flagellin and non-flagellin molecules. We hypothesized that Nlrc4 recognition of *Salmonella* flagellin is the dominant protective pathway during infection. To test this hypothesis, we used wild-type, flagellin-deficient, and flagellin-overproducing *Salmonella* to establish the role of flagellin in mediating Nlrc4-dependent host resistance during systemic and mucosal infection. We observed that during the systemic phase of infection, *Salmonella* efficiently evades Nlrc4-mediated innate immunity. However, during mucosal *Salmonella* infection, flagellin recognition by the Nlrc4 inflammasome pathway is the dominant mediator of protective innate immunity. These data establish that recognition of *Salmonella’s* flagellin by the Nlrc4 inflammasome during mucosal infection is the dominant innate protective pathway for host resistance against the enteric pathogen.

## Introduction

*Salmonella* is the causative agent in salmonellosis and is one of the main causes of gastrointestinal bacterial infections worldwide. Consumption of contaminated food is responsible for the majority of *Salmonella* infections and is the leading cause of foodborne-related deaths in the USA.^1^ During the initial phase of the infection, the bacteria travels to the intestine where it encounters a protective layer of mucus lining the intestines.^2^ After breaking through the mucus layer, *Salmonella* infects the intestinal epithelium and lamina propria myeloid cells, which are capable of detecting the bacteria through innate pattern recognition receptors (PRRs).^3-6^ PRRs recognize highly conserved structures of bacteria, such as rod and needle proteins from the *Salmonella* pathogenicity island 1 (SPI-1) type three secretion system (TTSS) and flagellin monomers that compose the flagellar filament.^7-14^

*Salmonella’s* flagellin, encoded by *fljB* and *fliC*, is a potent ligand that elicits a robust innate immune response by activating the caspase-1 (Casp1)-dependent inflammasome.^14-16^ Inflammasome recognition of flagellin is dependent on the detection of the conserved site, located on the carboxyl terminus of the protein, by Naip5 and Naip6 (Naip5/6).^7, 8, 12^ Flagellin recognition by Naip5/6 leads to the formation of a multiprotein complex, resulting in the activation of the Nlrc4-Casp1-dependent inflammasome.^17^ It has also been shown that the adaptor molecule ASC (apoptotic speck protein containing a caspase recruitment domain; encoded by *Pycard*) can associate with the Nlrc4 inflammasome resulting in efficient production of IL-1β; however, ASC-independent Nlrc4-Casp1 inflammasome activation still results in cell death via pyroptosis.^18^ Flagellin-mediated Nlrc4-dependent activation results in biologically active IL-1β and IL-18, eicosanoids, and Gasdermin-D-mediated pyroptotic cell death.^19, 20^

The SPI-1 TTSS is a critical virulence factor that enables *Salmonella* to colonize and successfully replicate in the host’s intestinal epithelial cells (IECs).^21^ Described as a needle-like structure, the TTSS delivers critical effector proteins, which allows *Salmonella* to create a favorable environment for colonization within IECs.^22, 23^ To counteract these virulence factors, the host uses PRRs, Naip1 and Naip2 (Naip1/2), which recognize the needle and rod proteins, encoded by *prgI* and *prgJ*, both of which are required for the TTSS needle complex assembly.^8, 9, 12^ Similar to flagellin, needle and rod recognition leads to the formation of the Nlrc4-Casp1 inflammasome, resulting in Nlrc4-dependent IL-1β production and pyroptotic cell death.^8, 9, 12^

Nlrc4 inflammasome recognition of *Salmonella* by IECs and phagocytes has been extensively studied and shown to be critical for innate host protection. It has been demonstrated that *Salmonella*-triggered activation of the Nlrc4 inflammasome in IECs not only results in the production of cytokine, eicosanoids, and pyroptosis, but also leads to IECs rapid expulsion from the intestinal epithelium.^3, 6^ More recently, it has been illustrated that Naip recognition of *Salmonella’s* pathogen associated molecular patterns (PAMPs) is critical for restricting the dissemination of the enteric pathogen.^24^

To evade Nlrc4 recognition, *Salmonella’s* expression of flagellin is tightly regulated by PhoPQ and ClpXP sensors and FlgM, which silences flagellin production *in vivo*.^25-28^ Experimental evidence suggest that SPI-1 TTSS is required for the intracellular translocation of flagellin, similarly regulated by two-component sensors, and silenced during intracellular and systemic infection.^29^ We hypothesized that flagellin recognition by the Nlrc4 inflammasome is the dominant pathway for host protection during mucosal *Salmonella* infection, when both flagellin and SPI-1 are upregulated to promote intestinal colonization and infection. To test our hypothesis, we used an attenuated strain of *Salmonella* that is unable to repress flagellin production (Δ*flgM*), in combination with Nlrc4-deficient (*Nlrc4*^-/-^) mice. Our results reveal that during intraperitoneal (i.p.) infection, *Salmonella* successfully evades recognition by the Nlrc4 inflammasome. In contrast, during mucosal infection, *Salmonella* production and secretion of flagellin via the flagellar basal body, but not SPI-1 TTSS, is detected by the Nlrc4 inflammasome and protects the host against infection. These data establish that flagellin recognition by the Nlrc4 inflammasome is critical for innate mucosal protection against *Salmonella enterica* serovar Typhimurium (*S*. Typhimurium).

## Methods

### Ethics Statement

This study was carried out in strict accordance with the recommendations in the Guide for the Care and Use of Laboratory Animals of the National Institutes of Health. All protocols were approved by the Institutional Animal Care and Use Committee of the University of Washington (protocol: 4031-01, Mucosal Immunity)

### Bacterial strains

The experiments were performed using wild-type (WT; from Brad Cookson, University of Washington), flagellin-deficient (Δ*fliC/fljB;* from Brad Cookson University of Washington), flagellin overexpressing (Δ*flgM*; a gift from Kelly Hughes), SPI-1 deficient (ΔSPI-1; from Kelly Hughes), flagellin overexpressing and SPI-1 deficient (Δ*flgM* /SPI-1; ΔSPI-1 from Kelly Hughes), flagellin-deficient and lacking control of flagellin repression (Δ*flgM*/*fliC/fljB*), flagellin overexpressing and lacking the flagellar basal body (Δ*flgM/flgB*; Δ*flgB* from Kelly Hughes), and SPI-2 deficient (ΔSPI-2; from Kelly Hughes) *S*. Typhimurium strain SL1344. Δ*flgM* mutant from Kelly Hughes was then transferred into Δ*fliC/fljB*, ΔSPI-1, ΔSPI-2, Δ*flgB* SL1344 strains using P22 phage.^30^ The deletion of *flgM* was confirmed by PCR. Bacteria were grown in Luria broth (LB) at 37°C with aeration.

### Mouse infection

C57BL/6 mice were purchased from Jackson Labs and housed in our facilities at the University of Washington. *Caspase-1*^-/-^ x *caspase-11*^-/-^ (*Casp1/11*^-/-^), *Nlrc4*^-/-^, and *Nlrp3*^-/-^ (generated by Genetech) were bred in our specific pathogen free (SPF) animal facilities.^31^ Animals were housed under standard barrier conditions in individually ventilated cages. 8-14 week old mice were used for infections throughout this study. All oral infections were performed as previously described.^15^ In brief, one day before infection, food was withdrawn 4 h prior to oral administration of 20 mg of streptomycin.^32^ Food was replaced and 20 h after streptomycin treatment, food was withdrawn again for 4 h prior to orally infecting mice with 1000 colony forming units (CFUs) of *S*. Typhimurium. Food was replaced immediately after infection. The *Salmonella* inoculum was prepared by back-diluting an overnight culture 1:50 in LB + 50 μg/ml of streptomycin. After 4 h, the concentration of bacteria was measured and diluted in cold PBS to a concentration of 1×10^4^ *CFU/ml*, and CFU of the inoculum was verified by plating on LB agar plates with 50 μg/ml streptomycin. Five days post-infection, mice were sacrificed by CO_2_ asphyxiation, tissues (intestine, mesenteric lymph node (mLN), spleen, and liver) were promptly removed. Bacterial burden was assessed by weighing and homogenizing the tissues in PBS with 0.025% Triton X-100, and plating dilutions of the samples on MacConkey agar plates with streptomycin (50 μg/ml). Prior to homogenization, the ceca were scraped and blotted to remove fecal content. *Salmonella* inoculum for systemic infections were prepared as previously described and 1000 CFUs of *S*. Typhimurium were administered intraperitoneally. Five days post-infection, mice were sacrificed by CO_2_ asphyxiation, tissues (spleen and liver) were promptly removed and bacterial burden was assessed as previously described.

### Quantitative histologic assessment

Formalin-fixed tissue was embedded in paraffin using standard protocols. 4 μm thick sections were stained with hematoxylin and eosin using standard procedures. A blinded pathologist examined the slides and scored them according to the following criteria. Scores were assigned for changes to the cecum as follows: submucosal expansion - 0 = no significant change, 1 = <25% of the wall, 2 = 25-50% of the wall, 3 =>50% of the wall; mucosal neutrophilic infiltrate - 0 = no significant infiltrate, 1 = mild neutrophilic inflammation, 2 = moderate neutrophilic inflammation, 3 = severe neutrophilic inflammation; lymphoplasmacytosis - 0 = no significant infiltrate, 1= focal infiltrates (mild), 2= multifocal infiltrates (moderate), 3 = extensive infiltrates involving mucosa and submucosa (severe); goblet cells - 0 = >28/HPF, 1 = 11-28/HPF, 2 = 1-10/HPF, 3 = <1/HPF; epithelial integrity - 0 = no significant change, 1 = desquamation (notable shedding of epithelial cells into the lumen), 2 = erosion (loss of epithelium with retention of architecture), 3 = ulceration (destruction of lamina propria). Crypt loss was estimated by blinded pathologist as fraction of cecal epithelium devoid of crypts.

### Macrophage cytotoxicity assay

Thioglycollate-elicited peritoneal macrophages were plated in a 96-well plate at a concentration of 5 × 10^5^ macrophages/well in RPMI 1640 medium with L-glutamine, 10% fetal bovine serum. *S*. Typhimurium was grown overnight in LB medium and back-diluted the next day 1:50 in LB medium and grown for 3-4 h. The bacteria were centrifuged and the pellet resuspended to the final desired concentration. Macrophages were infected with the desired multiplicity of infection (MOI), centrifuged at 250 x g for 5 min, and the infection was allowed to progress for an hour. Gentamicin (50 µg/ml) was added after an hour to kill extracellular bacteria. After an additional hour, the supernatants were removed and cytotoxicity was measured using Cytotox 96 kit (Promega).

### Statistics

Significance was obtained by using the software GraphPad Prism (San Diego, CA). One-way ANOVA was used when comparing three groups or more, using the Dunn’s multiple comparisons test. Two-way ANOVA was used when comparing three groups or more at multiple time points, using the Tukey’s multiple comparisons test. Statistical analyses of survival curve was done using Log-Rank (Mantel Cox) test. In all graphs, significance was established and represented using the following system: * = p<0.05, ** = p<0.01, *** = p<0.001, **** = p<0.0001.

## Results

### *S*. Typhimurium requires flgM to evade inflammasome detection during i.p. infection

The bacterial protein flagellin can be detected by Naip5/6, which activates the Nlrc4-dependent inflammasome. To evade recognition, *Salmonella* downregulates flagellin expression during host invasion. We tested whether the Nlrc4 inflammasome contributes to the control of systemic i.p. infection. To identify the role of Nlrc4 in host resistance against *S*. Typhimurium *in vivo*, we used mice lacking Casp1/11 (*Casp1/11*^-/-^), Nlrc4 (*Nlrc4*^-/-^), and Nlrp3 (*Nlrp3*^-/-^). We observed that upon i.p. infection with *S*. Typhimurium SL1344 (WT), there was no difference in the bacterial burden in the spleens or livers of mice deficient in the examined inflammasome components compared to C57BL/6 (B6) controls (Fig. 1A). Similarly, there was no difference in bacterial burden in the spleens and livers between B6 and the inflammasome-deficient strains of mice infected with flagellin-deficient (Δ*fljB*/*fliC*) *S*. Typhimurium (Fig. 1B). These data establish that inflammasome recognition is not the primary mediator of innate immunity during i.p. *S*. Typhimurium infection, suggesting that *Salmonella* efficiently evades inflammasome detection of flagellin and other potential ligands during i.p. infection.

**Figure 1.**
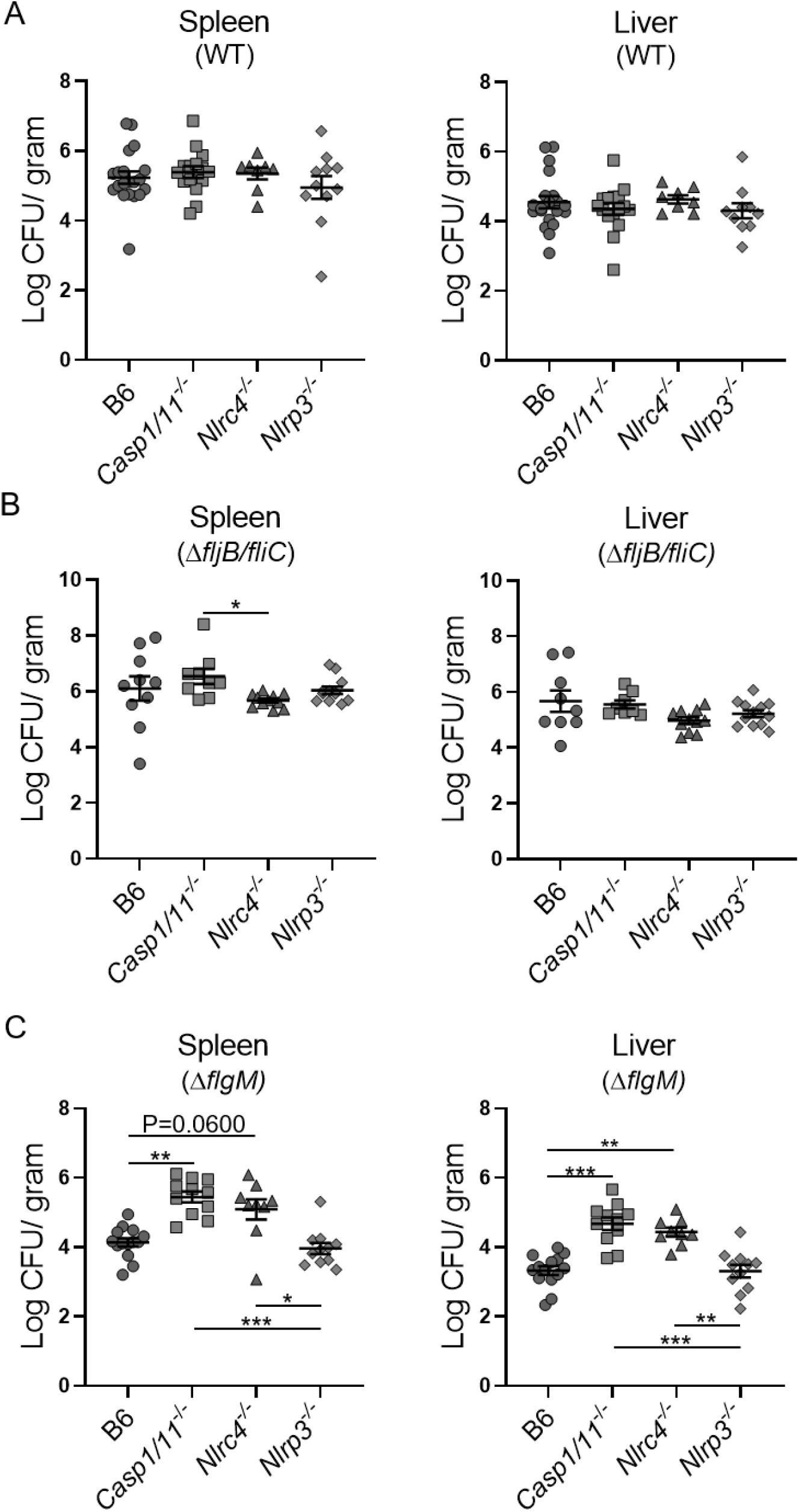
*S*. Typhimurium requires flgM to evade inflammasome detection during intraperitoneal infection. Bacterial burden of B6 (n=10-20), *Casp1/11*^-/-^ (n=9-15), *Nlrc4*^-/-^ (n=8-11), and *Nlrp3*^-/-^ (n=11-12) mice i.p. infected with 1000 CFUs WT SL1344 (**A**), Δ*fljB*/*fliC* (**B**), or Δ*flgM S*. Typhimurium (**C**) in the spleen and liver. Statistical analyses were done on using one-way ANOVA with Dunn’s multiple comparison test, *p<0.05, **p<0.01, ***p<0.001. Error bars, standard error mean.

*Salmonella* FlgM is an anti-sigma factor that binds FliA and prevents the expression of class III flagellar genes.^28, 33^ Upon completion of the flagellar basal body, FlgM is secreted and FliA is released to activate class III promoters, resulting in completion of the flagellar assembly. ^28, 33^ Deletion of *flgM* results in constitutive expression of flagellar class III genes and disruption of autogenous regulation of flagellar assembly.^28^ *flgM*-deficient (Δ*flgM*) *S*. Typhimurium expresses more flagellin protein, has more flagella than WT *Salmonella*, and is attenuated in mice.^15, 28^ We predicted that the attenuated phenotype observed in mice infected with Δ*flgM* S. Typhimurium is dependent on the response to flagellin by the Nlrc4 inflammasome. In striking contrast to both WT and Δ*fljB*/*fliC Salmonella* infections, Δ*flgM* i.p. infected *Nlrc4*^-/-^ and *Casp1/11*^-/-^ mice had dramatically elevated bacterial burden in both the spleen and liver compared to B6 and *Nlrp3*^-/-^ animals (Fig. 1C). Although Nlrp3 has been implicated in *Salmonella* detection during mucosal infection, we observed no phenotype for Nlrp3 in host protection against i.p. infection by WT, Δ*fljB*/*fliC*, or Δ*flgM Salmonella*.^34, 35^ Overall, these data indicate that the Nlrc4-Casp1/11 inflammasome is a critical mediator of flagellin detection; however, Nlrc4-Casp1/11-mediated immunity does not provide notable innate defense mechanism during i.p. infection of WT *Salmonella*.

### The Nlrc4-Casp1/11 inflammasome is critical for limiting mucosal inflammation and systemic *Salmonella* infection

lagellin is a key protein for bacterial motility; yet, *Salmonella* is capable of restricting flagellin expression based on its anatomical location in the host.^26^ Therefore, we investigated the role of the Nlrc4 inflammasome during oral infection in streptomycin pretreated mice. We observed rapid mortality in *Casp1/11*^-/-^ and *Nlrc4*^-/-^ mice compared to B6 controls orally infected with WT *S*. Typhimurium (Fig. 2A). Next, we measured the bacterial burden of *Casp1/11*^-/-^, *Nlrc4*^-/-^, and *Nlrp3*^-/-^ mice orally infected with WT *S*. Typhimurium. Our results showed no difference in the cecal bacterial burden between B6, *Casp1/11*^-/-^, or *Nlrc4*^-/-^ mice (Fig. 2B). We also observed that B6, *Casp1/11*^-/-^, and *Nlrc4*^-/-^ mice had elevated cecal bacterial burden compared to *Nlrp3*^-/-^ animals (Fig. 2B). Casp1/11- and Nlrc4-deficient animals had dramatically elevated bacterial burden in the mLN, spleen, and liver compared to B6 and *Nlrp3*^-/-^ mice (Fig. 2B). Infected *Nlrp3*^-/-^ mice had similar amounts of CFUs in the mLN, spleen, and liver compared to B6 animals (Fig. 2B). Histological examination revealed marked inflammation in all mice (Fig. 2C, D), and augmented tissue injury in mice lacking either Casp1/11 or Nlrc4 (Fig. 2E). These results reveal that the Nlrc4-Casp1/11 inflammasome plays a critical role in limiting intestinal tissue injury, as well as bacterial spread and growth in systemic sites during oral infections.

**Figure 2.**
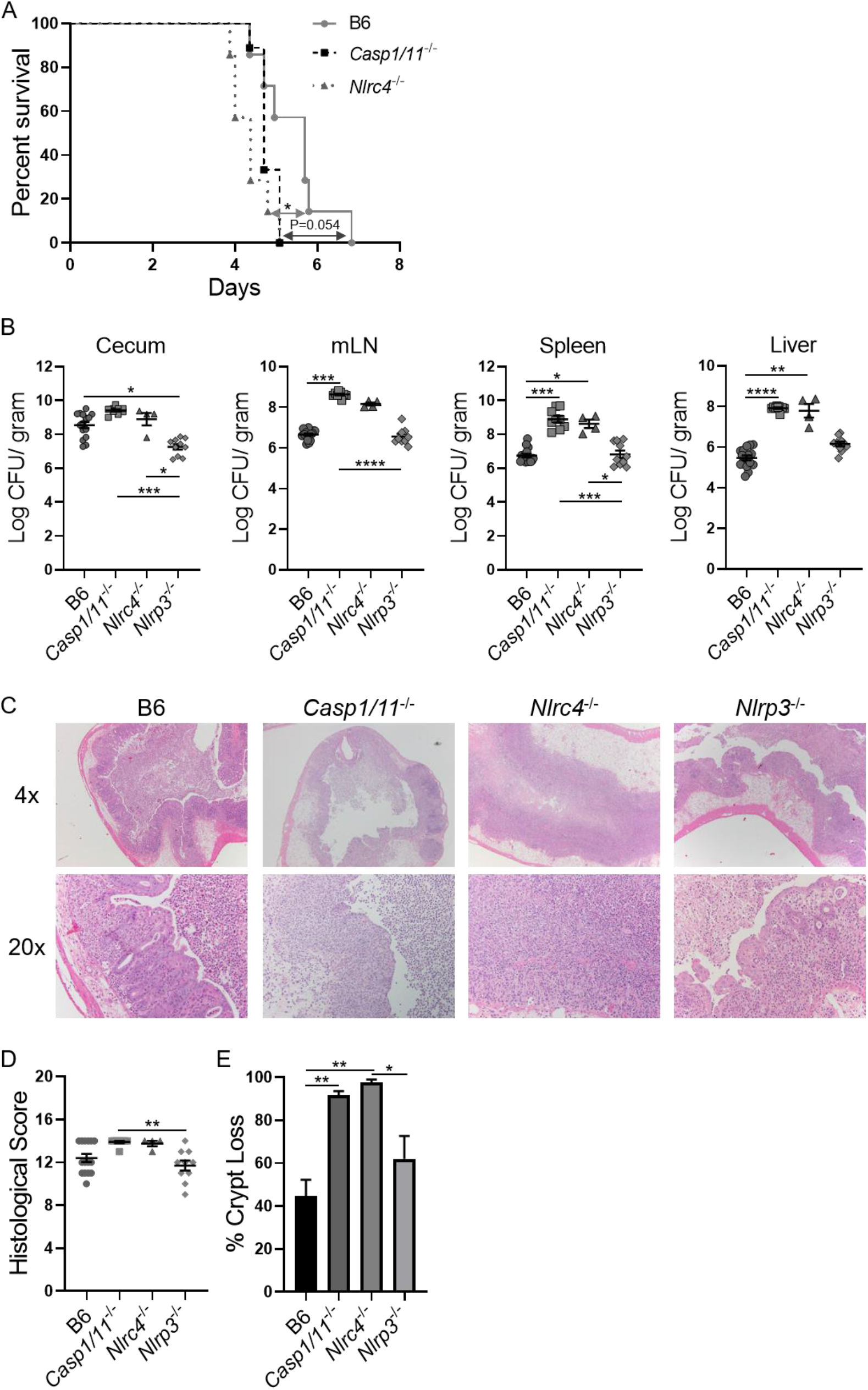
The Nlrc4-Casp1/11 inflammasome is critical for limiting systemic *Salmonella* infection. Survival of B6 (n=7), *Casp1/11*^-/-^ (n=9), and *Nlrc4*^-/-^ (n=7) mice that were orally infected with 1000 CFUs of WT SL1344 *S*. Typhimurium (**A**). Bacterial burden of B6 (n=15), *Casp1/11*^-/-^ (n=9), *Nlrc4*^-/-^ (n=4), and *Nlrp3*^-/-^ (n=10) mice orally infected with 1000 CFUs WT SL1344 *S*. Typhimurium in the cecum, mLN, spleen, and liver (**B**). Representative histology of the cecum infected with WT SL1344 *S*. Typhimurium (**C**). Histological scores for changes in the cecum (**D**). Frequency of crypt loss in the cecum (**E**). Survival curve is a combination of two-independent experiments involving at least 3 mice per group. Statistical analyses of survival curve was done using Log-Rank (Mantel Cox) Test. Statistical analyses were done on using one-way ANOVA with Dunn’s multiple comparison test, *p<0.05, **p<0.01, ***p<0.001, ****p<0.0001. Error bars, standard error mean.

### The Nlrc4 inflammasome protects against oral infection with *flgM*-deficient *Salmonella*

We have previously shown that Casp1/11 is critical for limiting the bacterial burden of mice orally infected with *flgM*-deficient *Salmonella*.^15^ Therefore, we hypothesized that the Nlrc4 inflammasome is also required for limiting growth of *Salmonella* lacking FlgM. To test our hypothesis, we orally infected inflammasome-deficient mice with Δ*flgM S*. Typhimurium. Our results show that in the absence of Casp1/11 or Nlrc4 there is a significant increase of *S*. Typhimurium in the cecum, mLN, spleen, and liver, compared to B6 and *Nlpr3*^-/-^ mice (Fig. 3A). Furthermore, Δ*flgM* orally infected *Casp1/11*^-/-^ and *Nlrc4*^-/-^ mice displayed extensive tissue destruction in the cecum (Fig. 3B-D). These data establish the Nlrc4-dependent inflammasome as essential for controlling bacterial burden and limiting intestinal pathology during infection.

**Figure 3.**
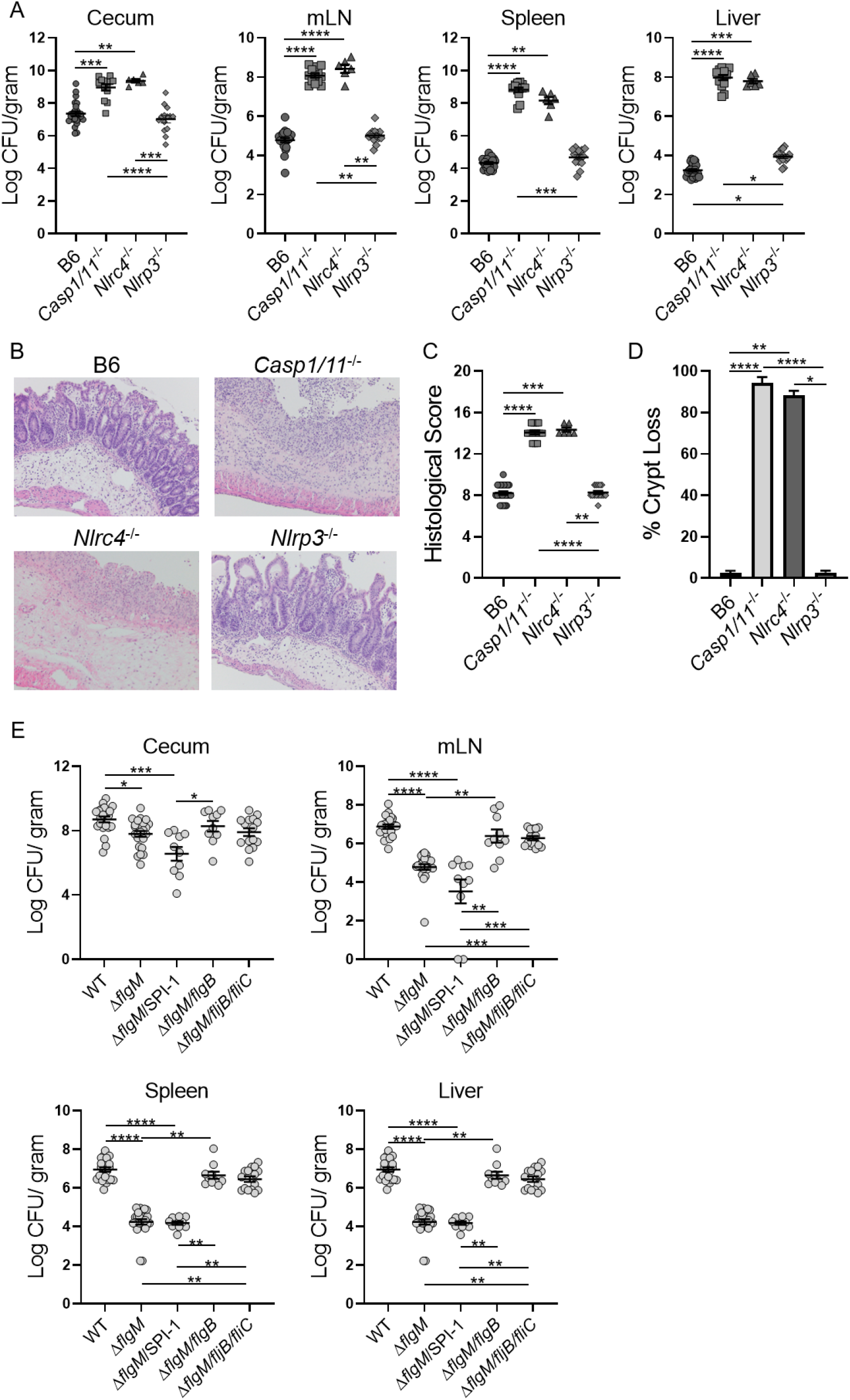
The Nlrc4-Casp1/11 inflammasome is required to mediate the attenuation of flgM-deficient *S*. Typhimurium. Bacterial burden of B6 (n=24), *Casp1/11*^-/-^ (n=14), *Nlrc4*^-/-^ (n=6), and *Nlrp3*^-/-^ (n=14) mice orally infected with 1000 CFUs Δ*flgM S*. Typhimurium in the cecum, mLN, spleen, and liver (**A**). Representative histology (20x) of the cecum infected with Δ*flgM S*. Typhimurium (**B**). Histological scores for changes in the cecum (**C**). Frequency of crypt loss in the cecum (**D**). Bacterial burden of B6 mice orally infected with 1000 CFUs of Δ*flgM* (n=24), Δ*flgM*/SPI-1 (n=10), Δ*flgM/flgB* (n=10), or Δ*flgM/fljB/fliC* (n=15) *S*. Typhimurium in the cecum, mLN, spleen, and liver (**E**). Statistical analyses were done on using one-way ANOVA with Dunn’s multiple comparison test, *p<0.05, **p<0.01, ***p<0.001, ****p<0.0001. Error bars, standard error mean.

### The flagellar basal body is required for Nlrc4 inflammasome detection of flagellin during mucosal infection

We next defined the requirement for *flgM*-deficient *Salmonella*-mediated inflammasome activation. To generate functional flagella, flagellin proteins are secreted through the flagellar basal body and polymerize into filaments.^33^ It has also been shown that flagellin activation of the inflammasome requires the SPI-1 TTSS, suggesting that flagellin is also secreted through SPI-1.^29^ To characterize both the flagellar basal body’s and SPI-1’s role in mediating flagellin-dependent *Salmonella* pathogenesis, we deleted the *flgB* gene or SPI-1. B6 mice were orally infected with WT, Δ*flgM*, Δ*flgM*/SPI-1, Δ*flgM*/*flgB*, or Δ*flgM*/*fljB/fliC Salmonella* and their bacterial burdens were assessed. We observed that the absence of both FlgM and SPI-1 had no effect on the pathogen burden compared to Δ*flgM* infected B6 mice (Fig. 3E). Conversely, in the absence of FlgM and the flagellar basal body, our results showed a significant increase of the bacterial burden in the mLN, spleen, and liver compared to Δ*flgM* and Δ*flgM*/SPI-1 infected mice (Fig. 3E). We also observed the attenuated phenotype of Δ*flgM Salmonella* was eliminated in the absence of flagellin expression (Fig. 3E). These results demonstrate that host recognition of *Salmonella* flagellin is primarily mediated by the protein secretion through the flagellar basal body and not SPI-1.

### Nlrc4 provides modest protection against aflagellate *Salmonella*

To determine the role of Nlrc4-inflammasome mediated detection of non-flagellin molecules in host resistance, we orally infected *Casp1/11*^-/-^ and *Nlrc4*^-/-^ mice with flagellin-deficient *Salmonella*. Compared to B6 mice, *Nlrc4*^-/-^ animals had significantly greater bacterial burden in the mLN, spleen, and liver (Fig. 4A). Although *Casp1/11*^-/-^ mice also displayed elevated bacterial burdens compared to B6 animals, this did not reach statistical significance (Fig. 4A). Histological analysis revealed no significant difference in inflammation between any strain of mice (Fig. 4B, C); however, *Nlrc4*^-/-^ and B6 mice did have slightly reduced tissue injury compared to Casp1/11-deficient animals (Fig. 4D). Overall, these data suggest that non-flagellin molecules are recognized by the Nlrc4 inflammasome pathway, but have only a limited role in restricting bacterial dissemination.

**Figure 4.**
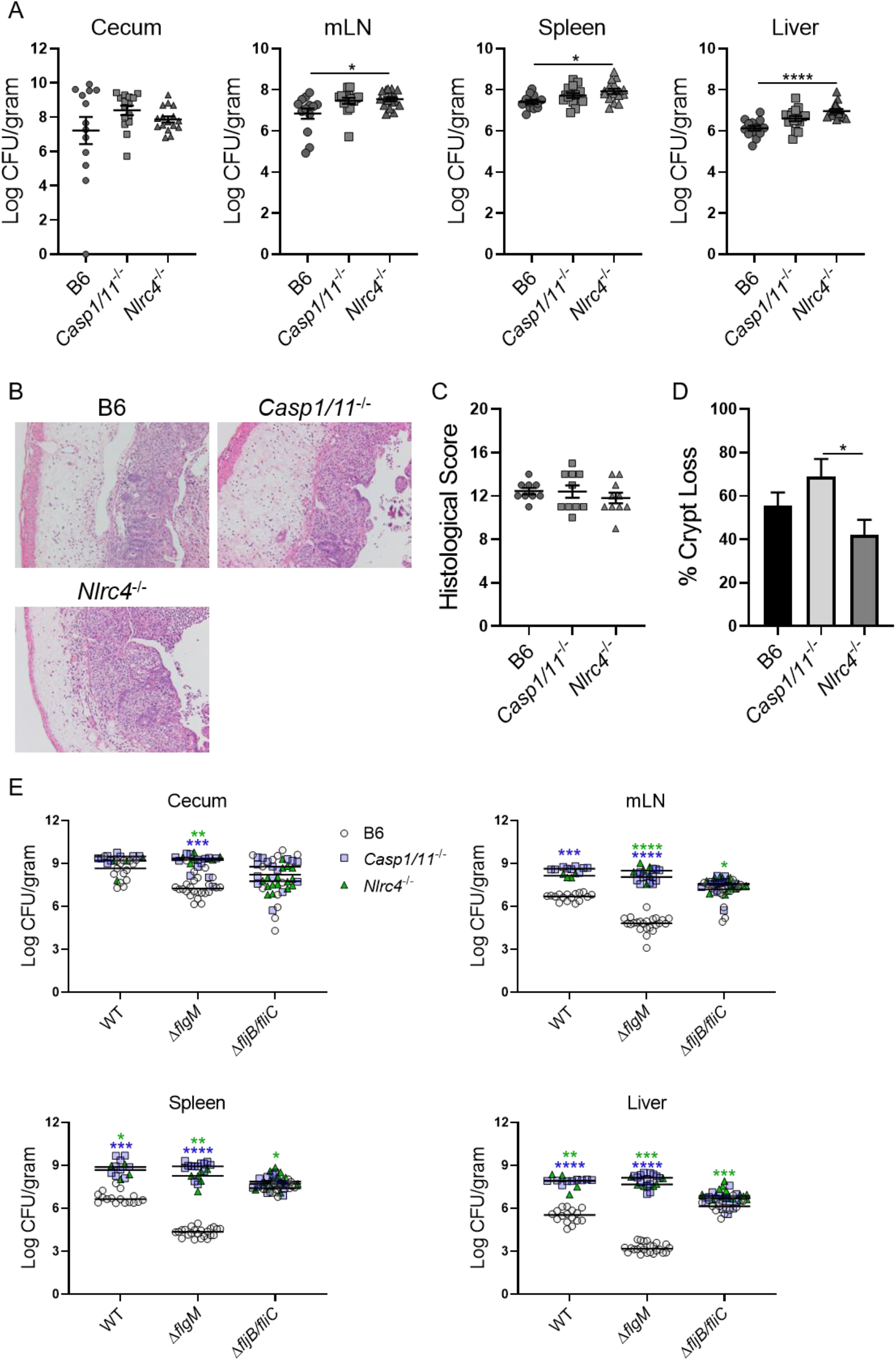
Nlrc4-Casp1/11-mediated host protection is dependent on flagellin expression. Bacterial burden of B6 (n=13), *Casp1/11*^-/-^ (n=15), and *Nlrc4*^-/-^ (n=16) mice orally infected with 1000 CFUs Δ*fljB/fliC S*. Typhimurium in the cecum, mLN, spleen, and liver (**A**). Representative histology (20x) of the cecum infected with Δ*fljB/fliC S*. Typhimurium (**B**). Histological scores for changes in the cecum (**C**). Frequency of crypt loss in the cecum (**D**). Composite analysis of WT SL1344, Δ*flgM*, and Δ*fljB*/*fliC S*. Typhimurium infections in B6, *Casp1/11*^-/-^, and *Nlrc4*^-/-^ mice (**E**). Statistical analyses were done on using one-way ANOVA with Dunn’s multiple comparison test, *p<0.05, **p<0.01, ***p<0.001, ****p<0.0001. Error bars, standard error mean.

To assess the overall changes in bacterial burden across all performed experiments, we compiled and compared CFU from B6, *Casp1/11*^-/-^, and *Nlrc4*^-/-^ mice orally infected with either WT, Δ*flgM*, or Δ*fljB*/*fliC Salmonella*. Our analyses demonstrate that the Nlrc4-Casp1/11 inflammasome is critical for the recognition of flagellin and significantly limits the bacterial burden in peripheral tissues such as the mLN, spleen, and liver (Fig. 4E). To a lesser extent, non-flagellin molecules recognized by the Nlrc4 inflammasome pathway also restrict bacterial growth in peripheral tissues. In addition, flagellin-dependent motility enhances the virulence of *S*. Typhimurium, which is also seen when comparing infection of *motA*-deficient *Salmonella* to WT and aflagellate *Salmonella* (Supplemental Fig. 1A). Our results suggest that during *Salmonella* infection, flagellin is the dominant ligand that is recognized by the Nlrc4 inflammasome pathway and required for efficient infection.

### Nlrc4-Casp1/11-mediated intestinal inflammation requires flagellin and SPI-1

To define the requirements for *Salmonella* efficiently activating the inflammasome, we tested *Salmonella* genes that are critical for oral infection in streptomycin pre-treated mice. *In vitro* Casp1/11-dependent killing of macrophages requires both flagellin and SPI-1 expression (Fig. 5A). Similarly, during oral infection in streptomycin treated mice, enhanced virulence of *Salmonella* in *Casp1/11*^-/-^ mice is dependent on flagellin and SPI-1 (Supplemental Fig. 2A). Augmented tissue inflammation of injury in *Casp1/11*^-/-^ mice relative to B6 mice was also dependent on SPI-1 and flagellin (Supplemental Fig. 1B), suggesting flagellin and SPI-1-dependent non-flagellin molecules are both required for enhanced virulence and to trigger intestinal pathology.

**Figure 5.**
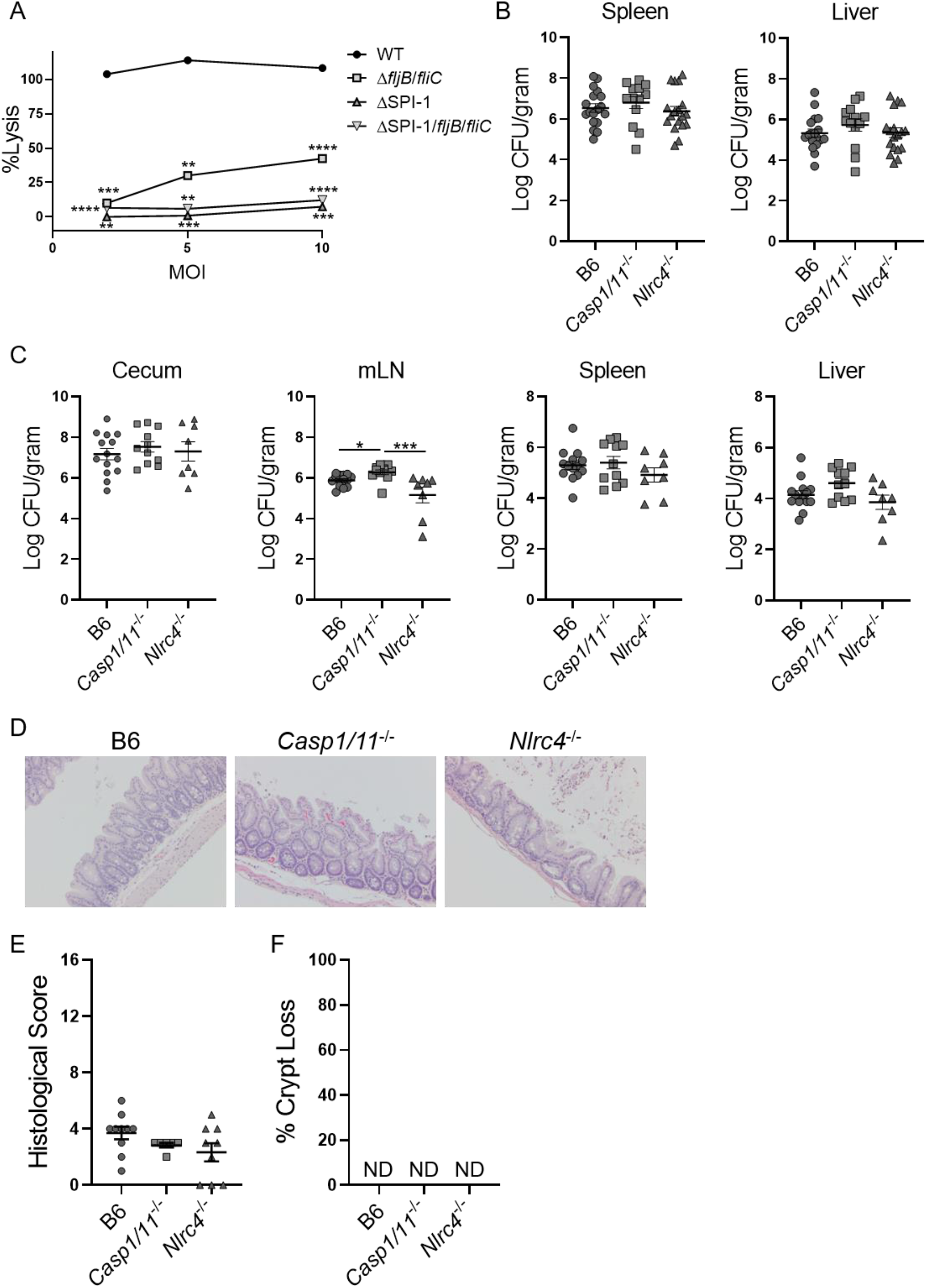
Flagellin-independent Nlcr4-Casp1/11-mediated intestinal inflammation is SPI-1-dependent. *Salmonella* induced cell death in thioglycollate elicited peritoneal macrophages measured by LDH release assay (**A**). Bacterial burden of B6 (n=18), *Casp1/11*^-/-^ (n=13), and *Nlrc4*^-/-^(n=18) mice i.p. infected with 1000 CFUs Δ*flgM*/SPI-1 *S*. Typhimurium in the spleen and liver (**B**). Bacterial burden of B6 (n=14), *Casp1/11*^-/-^ (n=11), and *Nlrc4*^-/-^ (n=8) mice orally infected with 1000 CFUs Δ*flgM*/SPI-1 *S*. Typhimurium in the cecum, mLN, spleen, and liver (**C**). Representative histology (20x) of the cecum infected with Δ*flgM*/SPI-1 *S*. Typhimurium (**D**). Histological scores for changes in the cecum (**E**). Frequency of crypt loss in the cecum (**F**). Statistical analyses were done using two-way ANOVA with Tukey’s multiple comparison test (A) or one-way ANOVA with Dunn’s multiple comparison test, *p<0.05, **p<0.01, ***p<0.001, ****p<0.0001 (B, C, E, F). Error bars, standard error mean; ND=not detected.

To assess the role of flagellin and SPI-1-dependent non-flagellin molecules in activating the Nlrc4 inflammasome, we infected mice with *Salmonella* lacking both flagellin and the entire SPI-1 needle complex (ΔSPI-1/*fljB*/*fliC*). We observed no differences between the bacterial burden of either i.p. or orally infected B6 and *Nlrc4*^-/-^ mice in all examined tissues (Fig. 5B, C). Yet, these results showed a subtle increase of CFUs in orally-infected *Casp1/11*^-/-^ compared to both B6 and *Nlrc4*^-/-^ animals in all tissues (Fig. 5C). Notably, examination of intestinal histology revealed that the absence of both flagellin and SPI-1 alleviated intestinal inflammation and tissue injury in all strains of mice (Fig. 5D-F). These data illustrate that flagellin and SPI-1 are required for efficient cecal colonization and *Salmonella*-induced intestinal inflammation and tissue injury. Overall, these results demonstrate that Nlrc4-mediated protection against *Salmonella* is dependent primarily on the expression of flagellin, and to a lesser extent SPI-1.

## Discussion

Previously, we demonstrated that flagellin recognition by Casp1/11 controls infection of *flgM*-deficient *S*. Typhimurium and limits intestinal inflammation and injury.^15^ It has been shown that *Salmonella*-mediated activation of the Nlrc4 inflammasome has distinct roles during systemic and mucosal infection through the detection of flagellin and non-flagellin molecules.^7-10, 12, 14^ In this article, we provide a more comprehensive understanding of how innate recognition of *S*. Typhimurium flagellin by the Nlrc4 inflammasome is essential for mucosal protection against the enteric pathogen. Notably, the inflammasome was not required for host defense against *Salmonella* infection when mice were infected via the i.p. route (Fig. 1A). This is likely due to the efficient downregulation of inflammasome ligands during the systemic phase of infection. In contrast, the Nlrc4 inflammasome is critical for prevention of *Salmonella*-induced intestinal tissue injury and systemic dissemination during mucosal infection (Fig. 2B). These data confirm that innate recognition of flagellin and SPI-1 TTSS structural proteins by the Nlrc4-Casp1/11 inflammasome is critical to limit bacterial burden and intestinal pathology during mucosal infections.

Previous studies have shown contradicting results as to the role of the Nlrp3 inflammasome in innate immunity against *S*. Typhimurium. The data reported by De Jong et al., and Hausmann et al., are consistent with our own, indicating a limited role for Nlrp3 in *Salmonella* resistance.^24, 36^ However, Broz and colleagues’ data indicate that innate recognition of *Salmonella* by the Nlrp3 inflammasome plays a significant albeit redundant role with Nlrc4 to limit *Salmonella* infection.^34^ Discrepancies between studies may be due to the limited number of mice tested and potential differences in gut microbiota that can influence oral infections. The preponderance of the data supports that for oral *Salmonella* infection in mice, the Nlrp3 inflammasome provides limited protection.

Using Δ*flgM S*. Typhimurium our data demonstrates that potent activation of the Nlrc4-Casp1/11 inflammasome pathway substantially limits bacterial burden and intestinal tissue damage (Fig. 3A-D). Because it has been shown *in vitro* that secretion of flagellin through the SPI-1 TTSS activates the inflammasome, we tested if the SPI-1 secretion pathway is required to activate the inflammasome *in vivo*. Our results establish that during oral infection with *flgM*-deficient *Salmonella*, FlgB-dependent secretion and assembly of flagella are required for flagellin-dependent activation of the inflammasome and that SPI-1 is not (Fig. 3E). Leakage of flagellin out of damaged *Salmonella*-containing vacuoles or escape of *Salmonella* into the cytosol are possible mechanisms for cytosolic delivery of flagellin that may be more relevant *in vivo*.

The critical role for Nlrc4 during mucosal *Salmonella* infection most likely reflects the need for flagella and the SPI-1 TTSS to efficiently invade host IECs. During oral WT *Salmonella* infection, SPI-1 and flagellin are both targeted by the Nlrc4 inflammasome. Using Δ*flgM S*. Typhimurium, our data indicates that recognition of flagellin by the Nlrc4-Casp1/11 inflammasome significantly reduces intestinal pathology and tissue injury (Fig. 3); likewise, intestinal tissue damage is augmented by deleting flagellin in *Salmonella* to levels seen in *Nlrc4*^-/-^ or *Casp1/11*^-/-^ mice (Fig. 4). SPI-1 is required to induce maximal intestinal inflammation and injury, and this is independent of the Casp1/11-inflammasome (Supplemental Fig. 2B). These results indicate that flagellin and SPI-1 are critical triggers of intestinal inflammation and injury through inflammasome-dependent and -independent pathways. Deleting both flagellin and SPI-1 alleviated *Salmonella*-mediated intestinal pathology and tissue damage, but did not prevent systemic dissemination of the bacteria (Fig. 5).

The limited phenotype for *S*. Typhimurium mutants lacking flagellin expression in B6 mice can be attributed to a concomitant loss of motility. This is most readily observed when looking at infection of B6 mice by Δ*motA S*. Typhimurium (Sup. Fig. 1A). When amotile flagellin-sufficient bacteria are compared to amotile flagellin-deficient bacteria, loss of flagellin expression results in enhanced virulence. Thus, motility enhances *Salmonella’s* virulence, which is offset by increased host resistance through the detection of flagellin by the Nlrc4-inflammasome. Nlrc4inflammasome induced inflammation also benefits *Salmonella* colonization of the gut and increases transmissibility, providing additional benefits for maintenance of flagellin-dependent motility in the face of host innate immune surveillance.

Since *Casp1/11*^-/-^ mice lack both canonical and non-canonical inflammasome pathways, the mucosal injury observed in these mice is independent of both the caspase-1 and caspase-11 inflammasomes. Epithelial cell intrinsic Naip-Nlrc4-Casp1 activation has been shown to induce cellular expulsion of infected enterocytes into the intestinal lumen, preventing S. Typhimurium infection of lamina propria mononuclear phagocytes, thereby restricting *Salmonella* dissemination to systemic tissues.^3^ We have previously shown that enhanced IL-12 and IFN-γ production by lamina propria leukocytes correlates with tissue injury.^15^ In addition, *Salmonella* accumulates more readily in lamina propria macrophages in the absence of Casp1/11.^15^ Thus, deficiency in the Nlrc4-Casp1 inflammasome may promote the accumulation of *Salmonella* within lamina propria macrophages and IFN-γ-dependent immunopathology.

Our study demonstrates that SPI-1 and flagellin are critical for efficient mucosal infection and the Nlrc4-inflammasome targets these virulence pathways to limit mucosal infection, inflammation, and tissue injury. Overproduction of flagellin in Δ*flgM S*. Typhimurium prevents excessive tissue injury, inflammation, and systemic spread in a Nlrc4-Casp1/11-dependent manner. Constitutive production of flagellin is a strategy to attenuate *Salmonella* while preserving the expression of this important virulence factor and target of innate and adaptive immunity. This may be useful for the development of live attenuated vaccines. Similar strategies to create *Salmonella* with constitutive SPI-1 expression may behave similarly and provoke protective innate immunity while maintaining the expression of important targets for adaptive immune responses.

## Figure Legends

**Supplemental Fig. 1.**
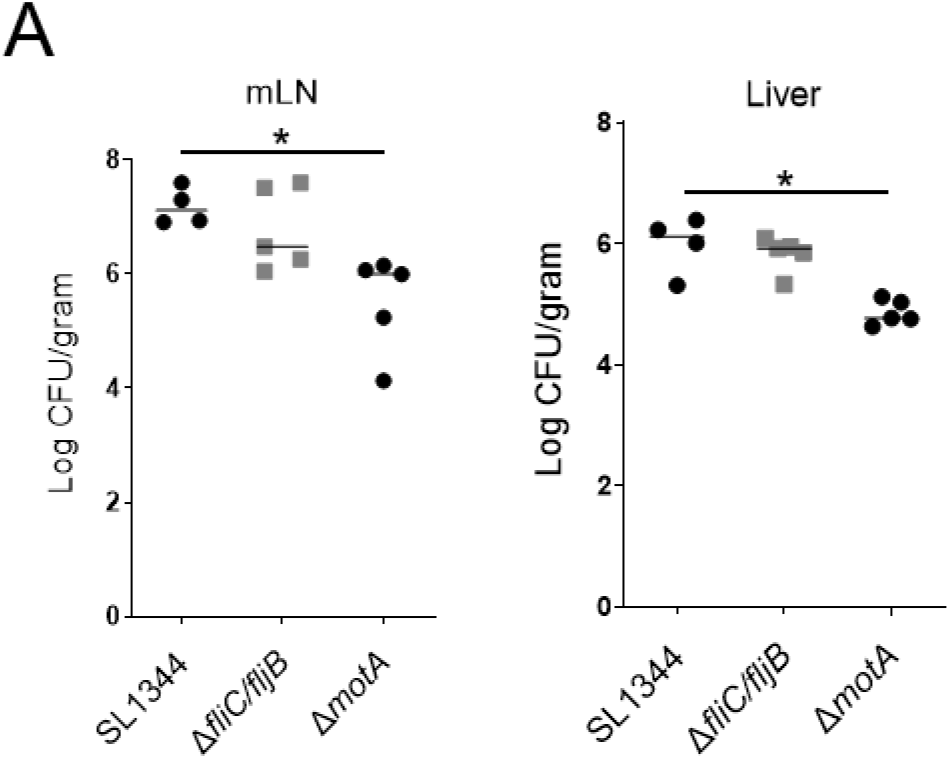
Bacterial burden of B6 mice orally infected with 1000 CFUs of WT SL1344, Δ*fljB*/*fliC*, or Δ*motA S*. Typhimurium in the mLN and liver (**A**). Statistical analyses were done one-way ANOVA with Dunn’s multiple comparison test, *p<0.05. Error bars, standard error mean.

**Supplemental Fig. 2.**
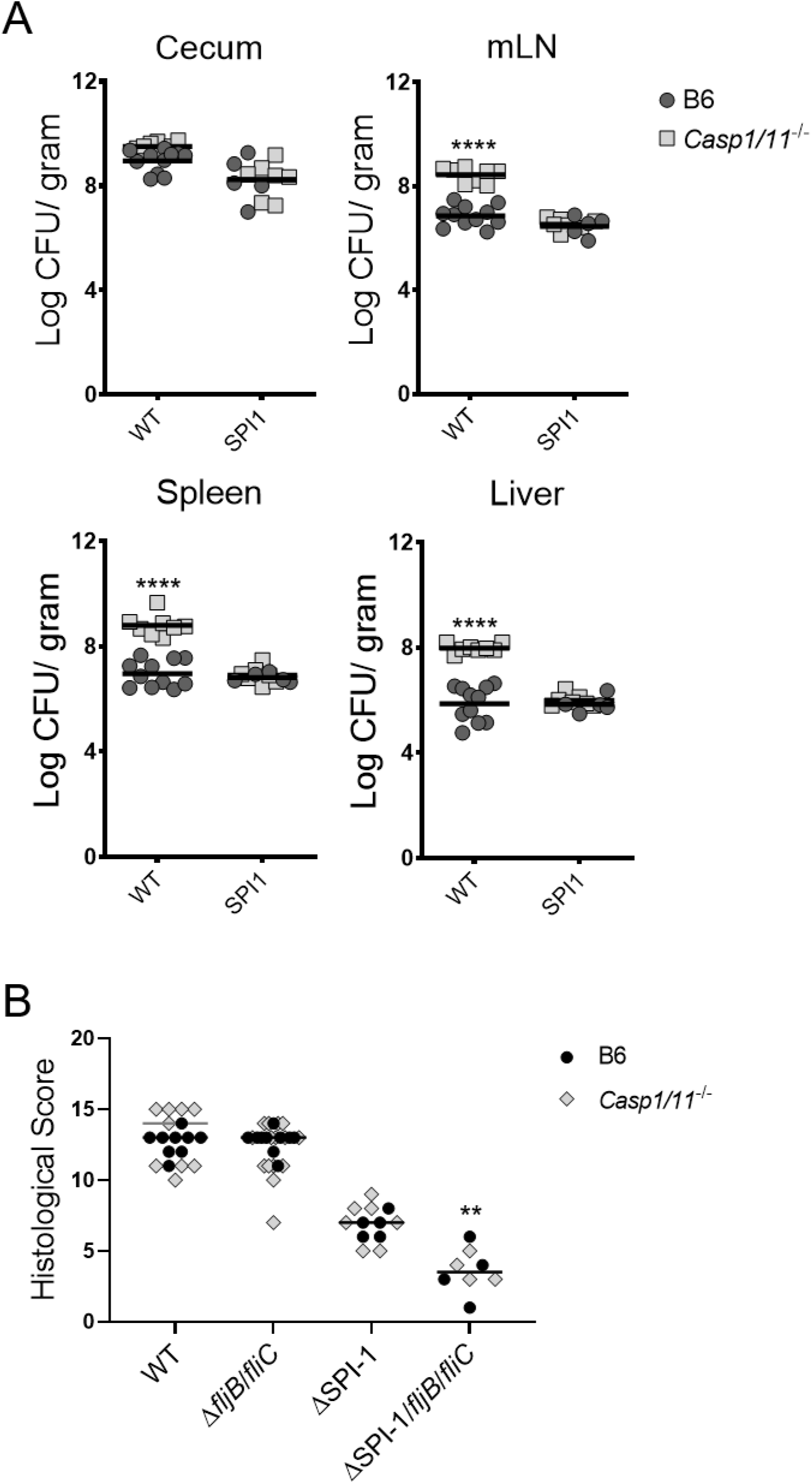
Bacterial burden of B6 and *Casp1/11*^-/-^ mice orally infected with 1000 CFUs of ΔSPI-1 *S*. Typhimurium in the cecum, mLN, spleen, and liver (**A**). Histological scores for changes in the cecum of B6 and *Casp1/11*^-/-^ mice orally infected with WT SL1344, Δ*fljB*/*fliC*, ΔSPI-1, ΔSPI-1/*fljB*/*fliC*, ΔSPI-2 *S*. Typhimurium (**B**). Statistical analyses were done with Mann-Whitney test, **p<0.01, ****p<0.0001. Error bars, standard error mean.

